# Whole-transcriptome causal network inference with genomic and transcriptomic data

**DOI:** 10.1101/213371

**Authors:** Lingfei Wang, Tom Michoel

**Affiliations:** Division of Genetics and Genomics, The Roslin Institute, The University of Edinburgh, Midlothian EH25 9RG, Scotland, United Kingdom

**Keywords:** causal gene network, whole-transcriptome network, causal inference, genome-transcriptome variation

## Abstract

Reconstruction of causal gene networks can distinguish regulators from targets and reduce false positives by integrating genetic variations. Its recent developments in speed and accuracy have enabled whole-transcriptome causal network inference on a personal computer. Here we demonstrate this technique with program Findr on 3,000 genes from the Geuvadis dataset. Subsequent analysis reveals major hub genes in the reconstructed network.

## 1 Introduction

Rapid developments in sequencing technologies have driven low the cost and high the throughput ([1]), with genomic and transcriptomic datasets from the same individuals increasingly publicly available (e.g. [2, 3]). The question now lies at the computational aspect, on how to fully exploit those datasets in order to address important biological and medical questions ([4]). Network-based approaches have received strong interests, especially from the clinical domain, where disease-related hub genes present attractive candidates for drug targeting ([5, 6]).

In this chapter, we focus on the reconstruction of causal gene networks on genome and (whole-)transcriptome datasets. (See [7] for a review.) As opposed to co-expression networks, causal gene networks are directed, and can identify the regulator among two co-expressed genes, or the existence of a hidden confounder gene or a feedback loop. With more stringent statistical tests, causal inference can also reduce the notoriously high numbers of false positives in co-expression networks.

For this purpose, genomic variations, which are typically observed in cohort studies and recombinant inbred lines, can be integrated as causal anchors or instrumental variables. Similar to double-blinded randomized controlled trials, genomic variations define naturally randomized groupings of individuals that allow to infer the causal relations between quantitative traits, a principle also known as Mendelian randomization ([8]). To test for a causal relation from a candidate regulator to a target, Mendelian randomization seeks a shared upstream causal anchor that is associated to both. By assuming that the causal anchor can only affect the target through the regulator, the interaction would then be identified.

However, this assumption does not always hold for gene regulations. For example, even if genomic variations are limited to lie in the cis-regulatory region of the regulator (i.e. cis-expression quantitative trait loci; cis-eQTLs), they may still also be associated to other nearby genes, which in turn control the target. Consequently, existing studies and public softwares, namely Trigger ([9]) and CIT ([10]), proposed to test this assumption through a “conditional independence test”. As was revealed in other studies, the conditional independence test cannot consider the existences of hidden confounders and technical variations ([11, 7, 12, 13, 14, 15]), which led to few discoveries of gene regulations. Additionally, neither software was efficient enough to handle the scale of modern datasets.

Recently in [14], we proposed alternative tests which are robust against confounders and technical variations. Implementational and statistical advances in the accompanying program Findr also resulted in almost 1,000,000 times of speed-up compared to CIT. This makes possible the reconstruction of whole-transcriptome causal gene networks, which can detect novel interactions by avoiding any pre-selection of genes.

In this chapter, we present a detailed protocol for the application of Findr, through an example where causal gene networks are inferred among 3000 genes from down-sampled Geuvadis study data ([2]). This is supplemented with a brief outline of the methods implemented in Findr, and its future perspectives in method development and application domains.

## 2 Notations and materials

In this section, we briefly formalize the network inference question and establish the necessary computational environment for Findr.

### 2.1 Question formalization

Consider genome-transcriptome variation data from *N* unrelated individuals. After preprocessing and eQTL analysis, we have identified *G* expressed genes (*see* Note 1), *g*_1_*,g*_2_*,…,g*_*G*_, in which the first *E ≤ G* genes (i.e. *g*_1_*,g*_2_*,…,g*_*E*_) have cis-eQTLs.

We denote the expression level (FPKM) of gene *g*_*i*_ for individual *j* as *g*_*i,j*_, whose matrix form is

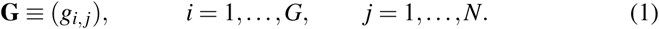

Similarly, for gene *g*_*i*_, the genotype of its cis-eQTL for individual *j* is defined as *e*_*i,j*_, with the matrix form

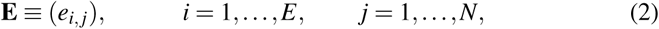

where each genotype is limited by the number of alleles, *N*_*a*_, as

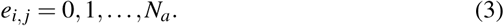

The unknown gene regulation network can be represented as the posterior probability of regulation between every pair of genes, given the observed data, as

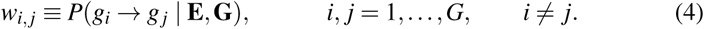

Regulations are identified solely according to their expression and eQTL patterns, independent of the underlying mechanism or whether the regulation is direct (*see* Note 2).

Causal inference utilizes the cis-eQTL of every regulator gene to map the probability of regulation for all its possible targets. Therefore, genes without any cis-eQTL (*g*_*E*+1_*,…,g*_*G*_) are regarded as only target genes but not as regulators, with

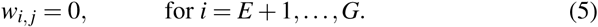

The expression levels of all possible regulators is also a sub-matrix of **G** as

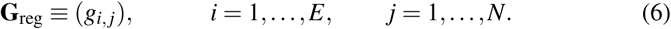

Given the expression levels **G** and cis-eQTL genotypes **E**, the question is to compute the probability of regulation matrix:

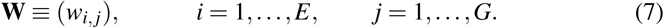

### 2.2 The Findr program

Findr (Fast Inference of Networks from Directed Regulations) is an efficient C library to reconstruct causal gene networks, whose methods can deal with the unique challenges in genomic and transcriptomic datasets, and are sketched in Sect. 4. Findr provides interfaces in python2, R, and command line (*see* Note 3). Its efficient implementation and analytical calculation of null distributions (Sect. 4.2.2) are pivotal to the speed-up of nearly 1 million times compared to existing programs [14]. This allows for whole-transcriptome causal network inference on modern datasets.

As a demonstrative example, we use the Findr R package with version 1.0.3 ([16]) in this chapter (*see* Note 4).

### 2.3 Computing environment

At the time of writing, the latest Findr R package (version 1.0.3) requires the following computing environment:

- A modern personal computer, or a high performance computing environment (*see* Note 5).
- A modern Linux or Mac operating system.
- The GCC compiler (*see* Note 6).
- A command-line environment (to install Findr).
- A recent R language environment (R, RStudio, etc).

## 3 Whole-transcriptome causal network from the Geuvadis dataset

The Geuvadis project [2] measured genome-wide genotypes and gene expression levels in 465 human lymphoblastoid cell line samples. Using this dataset as an example, here we reconstruct a causal gene network with Findr in R.

### 3.1 Install Findr

The latest version of Findr (*see* Note 7) can be downloaded and installed with the following lines in command-line environment (*see* Note 8):

#Comments above a command explains its function

#Comments below a command (if present) shows its expected output

#Download Findr R package from github (*see* Note 9)
git clone https://github.com/lingfeiwang/findr-R.git

#Install Findr
cd findr-R && R CMD INSTALL findr

### 3.2 Prepare data

Here we reconstruct a causal gene regulation network among 3000 genes in which 1000 have cis-eQTLs. The dataset was downsampled from the Geuvadis project (*see* Note 10). In R, the Findr library and the downsampled Geuvadis dataset can be loaded with:

#Load Findr
library(findr)

#Load downsampled Geuvadis dataset
data(geuvadis)

### 3.3 Reconstruct network

Network inference is performed with the function *findr.pij gassist*, taking **E**, **G**_reg_, and **G** as input and returning **W** as ouput (*see* Note 11):

#Reconstruct causal gene network
w=findr.pij gassist(geuvadis$dgt,geuvadis$dt,geuvadis$dt2,nodiag=TRUE)

#Examine output dimension
print(dim(w))

#[1] 1000 3000

The computation takes about one second on a modern desktop computer, and scales linearly with the numbers of regulators, targets, and individuals.

### 3.4 Analyze and visualize network

To demonstrate the properties of the reconstructed causal network for human lymphoblastoid cell lines, we briefly analyze and visualize it below:

#### 3.4.1 Regulation probabilities

The distribution of posterior regulation probability *P*(*g*_*i*_ → *g*_*j*_ | **E,G**) is visualized in Fig. 1 with code (*see* Note 12):

**Fig. 1.**
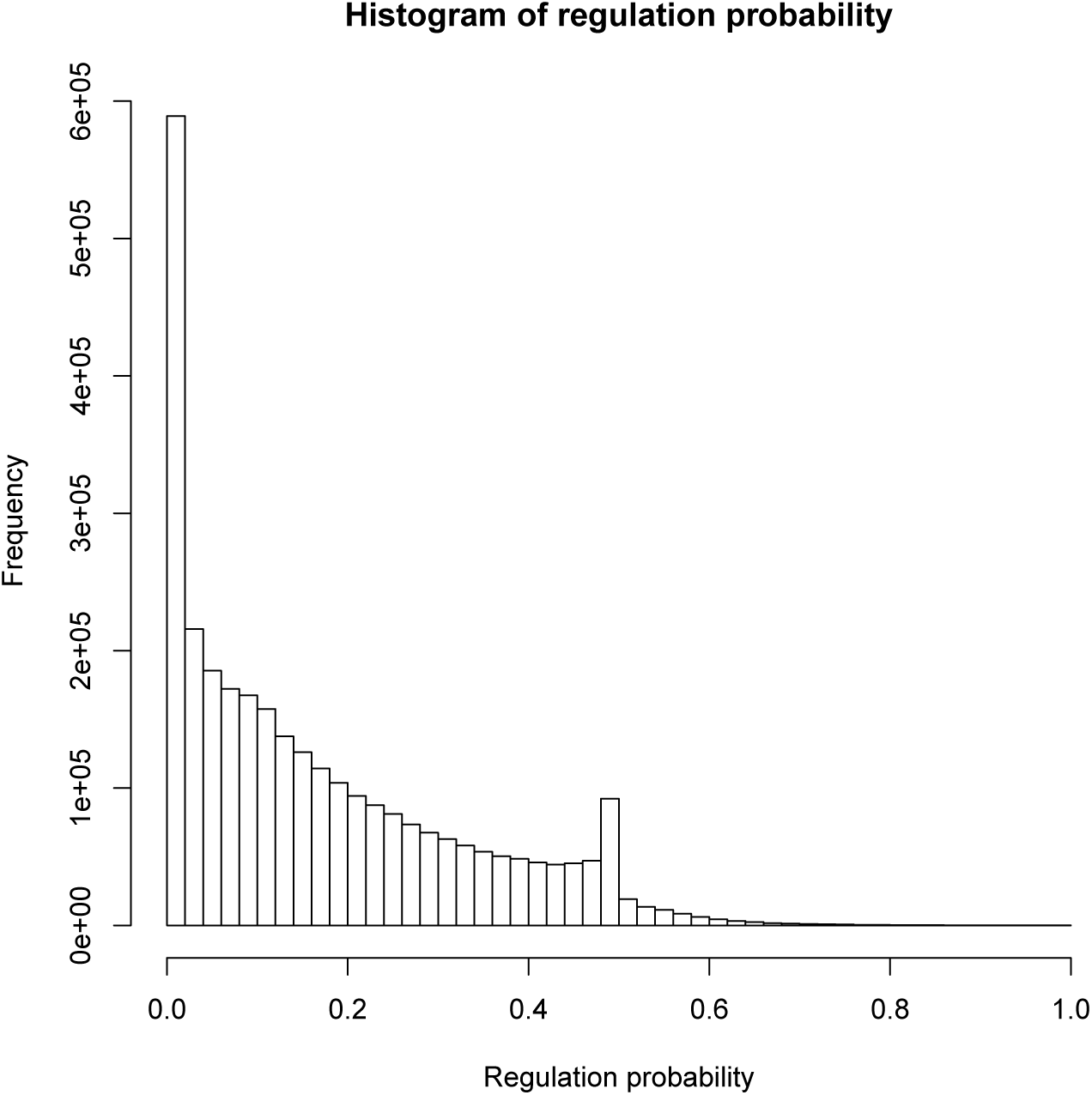
The histogram of regulation probability (off-diagonal elements of **W**) of the causal gene network reconstructed from the downsampled Geuvadis dataset.

#Drop self-regulation probabilities
wnd=w
diag(wnd)=NA
wnd=w[which(!is.na(wnd))]

#Histogram of regulation probabilities
hist(wnd,breaks=50,main=‘Histogram of regulation probability’,
xlab=‘Regulation probability’)

#### 3.4.2 Out-degree distribution

The distribution of out-degree is visualized in Fig. 2 with code:

**Fig. 2.**
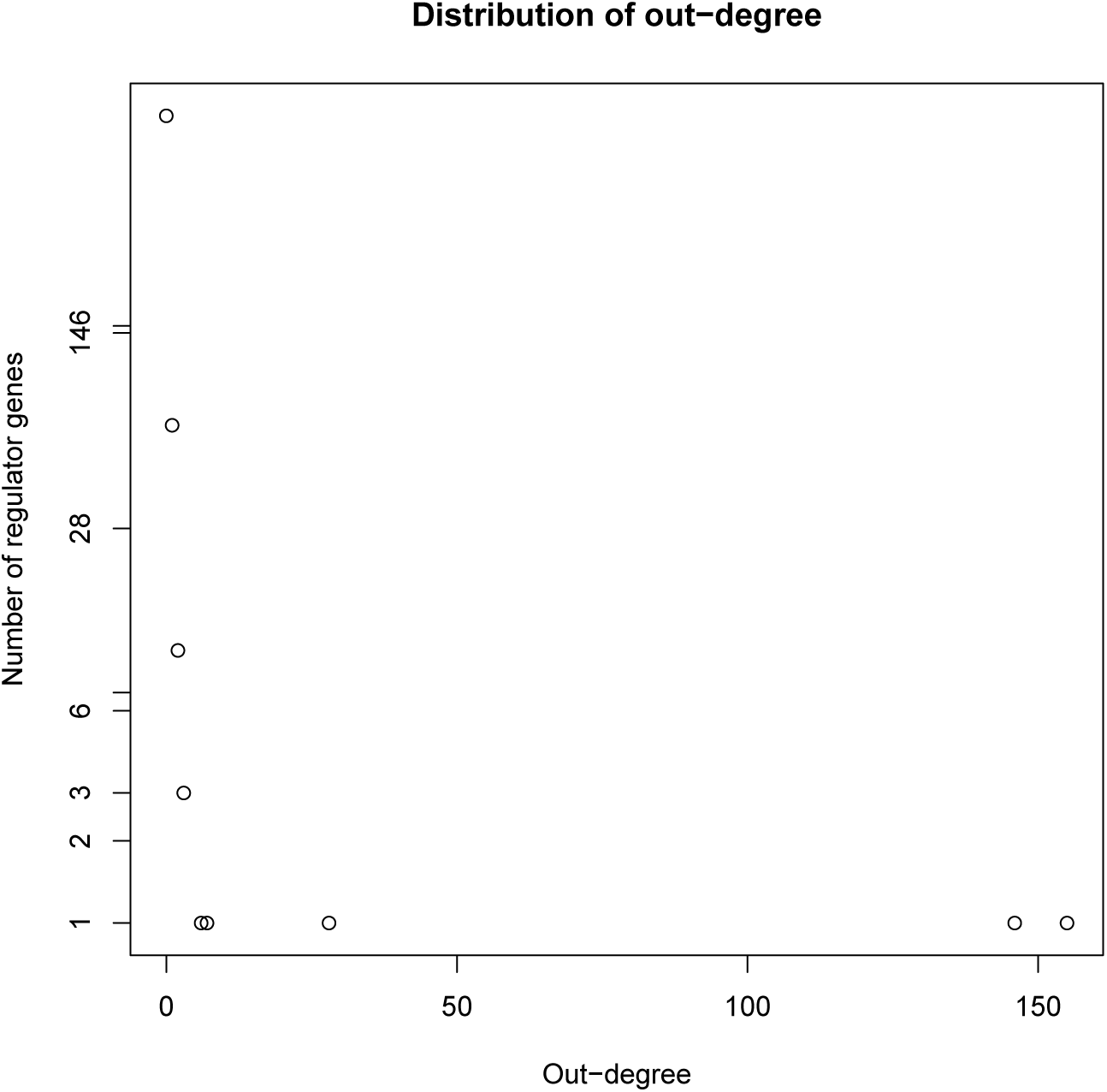
The histogram of out-degree of regulator genes in the thresholded causal gene network reconstructed from the downsampled Geuvadis dataset.

#Threshold network to 90% confidence level
threshold=0.9
net=w*>*=threshold

#Histogram of out-degree
odeg=rowSums(net)
vals=table(odeg)
plot(names(vals),vals,log=‘y’,main=‘Distribution of
out-degree’,xlab=‘Out-degree’,ylab=‘Number of regulator genes’)

#### 3.4.3 Top hub gene list

The list of top hub genes (by out-degree) is obtained below:

#Find top 5 hub genes, and their numbers of targets
ntop=5
odego=odeg[order(odeg,decreasing=TRUE)]
print(odego[1:ntop])

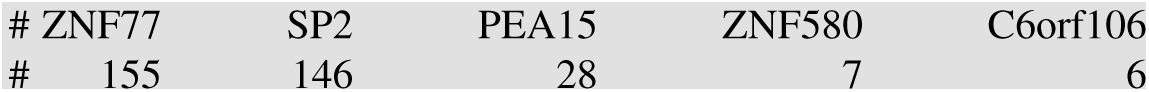

#### 3.4.4 Network visualization in Cytoscape

The reconstructed network can be exported to a csv file for visualization in Cytoscape (*see* Note 13), with the following code:

#Convert causal network to sparse format
dat=NULL
for (i in 1:dim(net)[1])
if(any(net[i,]))
dat=cbind(dat,rbind(rownames(net)[i],names(which(net[i,]))))
dat=t(dat)
colnames(dat)=c(‘source’,‘target’)
print(dat[1:2,])

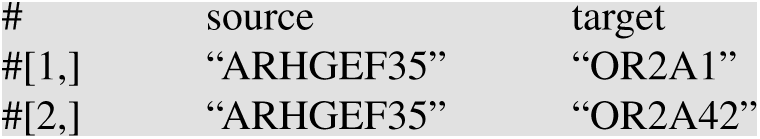

#Export sparse network to dat.csv
write.csv(dat,‘dat.csv’,row.names=FALSE,quote=FALSE)

Cytoscape can then import the network in dat.csv and visualize it. The largest connected component is shown in Fig. 3.

**Fig. 3.**
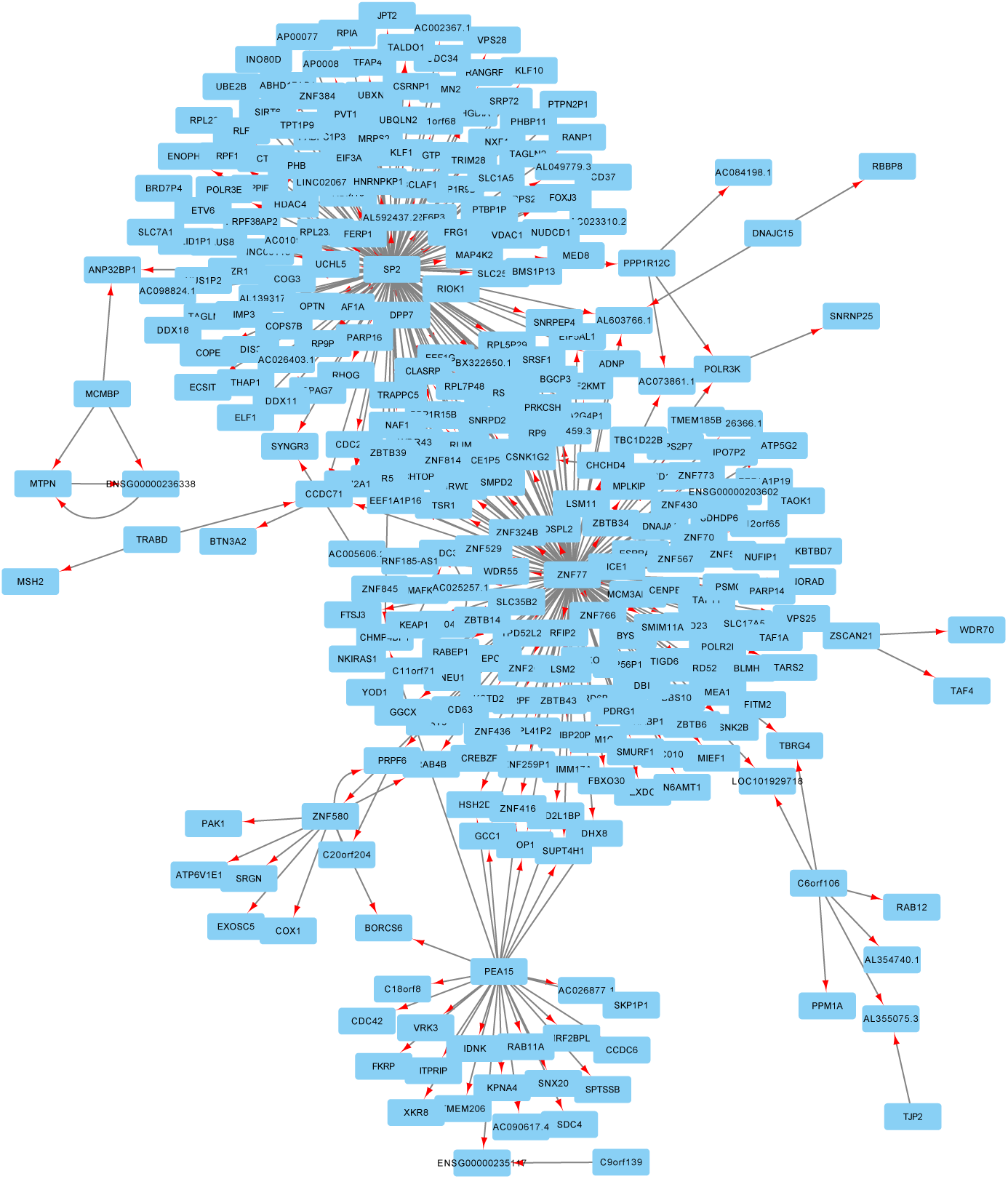
The largest connected component of the reconstructed and thresholded regulation network from the downsampled Geuvadis dataset, as visualized in Cytoscape. Of the 3000 genes, 308 are in this connected component with 366 regulations.

## 4 Statistical Methods

In this section, we outline the statistical methods used in Findr. (For details, see
[14].)

### 4.1 Data normalization

To satisfy the assumptions of linear dependency and normal noise distribution, and also to remove outliers (*see* Note 14), the expression levels of each gene are transformed to follow the standard normal distribution, based on the expression level ranking across individuals. Each gene is normalized separately.

### 4.2 Causal inference subtests

Consider all possible regulatory relations between the triplet (*A,B,C*), in which *B* is the regulator gene, *A* is its cis-eQTL, and *C* is a potential target gene. Causal inference performs three subtests in Table 1 (*see* Note 15) to narrow down their relation to contain *A → B → C*, but with a false positive rate as low as possible. Each subtest compares a null and an alternative hypothesis, by first performing a likelihood ratio test, then converting the likelihood ratio into p-values, and finally computing the posterior probability of alternative hypotheses given the observed data, which is equivalent with the *local* False Discovery Rate (FDR). This is similar with Genome Wide Association Studies, in which the Pearson correlation (equivalent with likelihood ratio) is first computed, and then converted into p-values and FDRs.

**Table 1.**
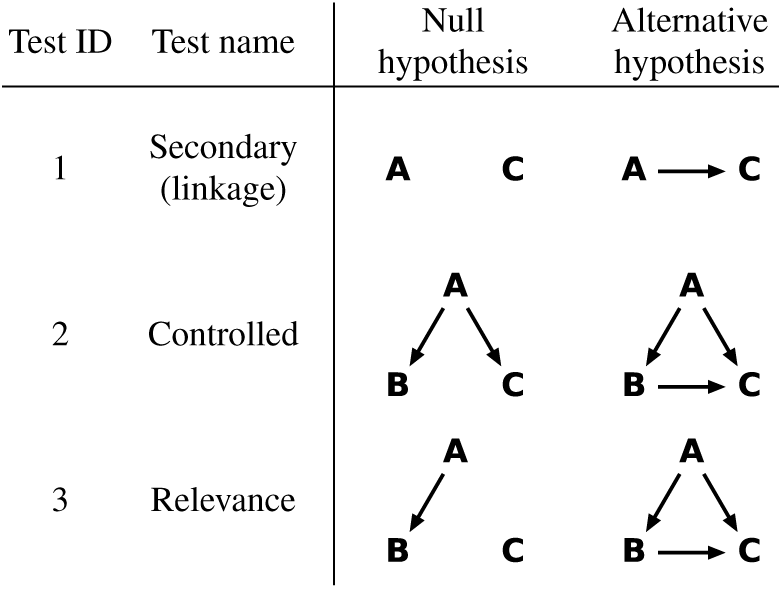
Three subtests of causal inference performed to test the gene regulation *B → C*, with the cis-eQTL of *B* as the causal anchor *A*. These tests are numbered, named, and defined in terms of null and alternative hypothese as shown. Arrows in a hypothesis indicate directed regulatory relations. (From [14].)

#### 4.2.1 Likelihood ratio tests

In Table 1, each graph represents a probablistic dependency model among the (*A,B,C*) triplet. The expression level of each gene is modelled as following a normal distribution, whose mean depends additively on all its parents. The dependency is linear on other gene expressions, and categorical on genotypes. Based on the normally distributed models, the likelihood ratio between the alternative and the null hypotheses can be computed for each subtest.

#### 4.2.2 P-values

For each subtest, the null distribution of likelihood ratio may be obtained either by simulation or analytically. Regardless of the method, likelihood ratios can then be converted into p-values according to their rankings in the null distribution. In [14], we found that the null distribution can be computed analytically, therefore avoiding simulations and accelerating the computations in Findr by *∼* 1000 times.

#### 4.2.3 Posterior probabilities of alternative hypotheses

P-values can be further reformulated into the posterior probabilities of alternative hypotheses, according to [17, 9]. As opposed to FDRs, the combination of subtests requires posterior probabilities which correspond to local FDRs (*see* Note 16). Findr implements a simplified estimator (*see* Note 17), with the resulting posterior probabilities denoted as

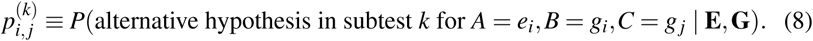

### 4.3 Subtest combination

Findr computes the final probability of regulation by combining the subtests as:

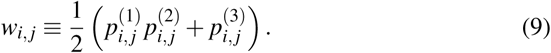

By combining the secondary linkage and controlled tests, the first term verifies that the correlation between *g*_*i*_ and *gj* is not entirely due to pleiotropy. By replacing the conditional independence test in [9] with the controlled test, this combination is robust against hidden confounders and technical variations.

On the other hand, the relevance test in the second term can identify interactions that arise from the indirect effect *e*_*i*_ → *g*_*i*_ → *g*_*j*_ but are too weak to be detected by the secondary linkage test. However, in such cases the direction of regulation cannot be determined. The coefficient 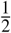 simply assigns half of the probability to each direction.

## 5 Notes

1. The whole chapter is equally applicable on gene isoforms.

2. However, direct regulations tend to have stronger significance than the indirect regulation they form, if the relevant genes have similar levels of technical variations.

3. URLs for Findr library and interfaces: - Library: https://github.com/lingfeiwang/findr - Command line interface: https://github.com/lingfeiwang/findr-bin - Python2 interface: https://github.com/lingfeiwang/findr-python - R package: https://github.com/lingfeiwang/findr-R

4. Findr’s python2 and command-line interfaces have the same functionality with similar computing environment requirements.

5. The computer memory (RAM) limits the size of data Findr can process. Findr’s python2 and command-line interfaces can automatically split the dataset according to the specified memory usage limit. The R package requires manual division. A modern computer with 4GB of RAM can handle whole-transcriptome network inference for hundreds of individuals, with thousands of genes having cis-eQTLs and tens of thousands having not.

6. GCC is already installed on most Linux machines. On Mac OS, Apple’s compiler may pretend to be GCC, so Findr may fail to install. For the solution, see FAQ in https://github.com/lingfeiwang/findr/blob/master/doc.pdf. GCC can be downloaded from https://gcc.gnu.org.

7. To exactly reproduce the example, Findr 1.0.3 can be downloaded from [16].

8. Detailed installation instructions and FAQs are available in the full manual at https://github.com/lingfeiwang/findr/blob/master/doc.pdf, or the doc.pdf file of the corresponding version.

9. Without git, the latest version of Findr can also be downloaded from https://github.com/lingfeiwang/findr-R/archive/master.zip. The user then needs to uncompress it and enter the corresponding folder before installation.

10. The original, full Geuvadis dataset can be downloaded from ArrayExpress (https://www.ebi.ac.uk/arrayexpress/), in accessions E-GEUV-1 and E-GEUV-2. In this analysis, 360 unique European individuals of 465 in total are considered. Geuvadis reported 23722 genes expressed (in ≥ 50% individuals) after QC. Genetic loci without any variation were discarded. After that, the Geuvadis QTL analysis identified 3172 genes with at least one significant cis-eQTL. Findr includes a random subset of these data for illustration purposes, totalling 1000 genes with cis-eQTLs and 2000 more without cis-eQTLs. For each of the 1000 genes, the genotypes of its strongest cis-eQTL are also included. The same network analysis can be performed on other datasets, such as on the full Geuvadis dataset to reconstruct the whole-transcriptome causal gene networks among 23722 genes. This requires an already performed eQTL analysis for the dataset, either from softwares such as matrixeQTL ([18]) or fastQTL ([19]), or from existing studies.

11. Function description for findr.pij gassist(dg,dt,dt2,na=NULL,nodiag=FALSE): - dg: Input integer matrix **E** of cis-eQTL genotype data, as defined in Eq 2. The element [*i, j*] is the genotype value (0,1*,…,N*_*a*_) of the cis-eQTL of regulator gene *i* for individual *j*. - dt: Input double matrix **G**_reg_ of regulator gene expression level data, as defined in Eq 6. The element [*i, j*] is the expression level of regulator *i* for individual *j*. - dt2: Input double matrix **G** of all gene expression level data, as defined in Eq 1. The element [*i, j*] is the expression level of gene *i* for individual *j*. - na: The number of alleles *N*_*a*_. If unspecified, it is automatically detected as the maximum value of dg. - nodiag: This function can infer networks for regulators that either should or should not be regarded as targets. In the earlier case, the regulators should also appear before other genes as targets, and nodiag should be TRUE. In the latter case, there should be no overlap between regulators and targets, with nodiag=FALSE. - Return value: Output double matrix **W** of inferred probability of regulation, as defined in Eq 7. The element [*i, j*] is the probability of regulation from regulator *i* to target *j* after observing the input data.

12. The peak at regulation probability around 0.5 is due to correlated regulator and target genes, whose regulation direction cannot be determined with the cis-eQTL of the regulator. In such cases, the novel combination of causal inference tests assumes a half probability for each direction.

13. http://www.cytoscape.org/

14. Although the data normalization step attempts to transform gene expression levels into the standard normal distribution, ill-distributed datasets may still cause underperformance from the method and should therefore be analyzed with extra care. Examples may include a large proportion of ties in gene expression levels, from single-cell transcriptomics or sparsely expressed genes.

15. Here we omit the primary linkage test, comparing *A → B* against *A B*, because the cis-eQTL *A* is assumed to be significant. These tests are also numbered differently with [14].

16. For the difference between FDR and local FDR, see [20].

17. Findr also skips the computation of P-values when deriving the posterior probabilities using the null distribution. For more stringent local FDR conversion with Grenander estimator, the user can first compute p-values within Findr (findr.pijs gassist pv) and then use other softwares (e.g. fdrtool in R from [20]) to obtain local FDRs.

## 6 Future perspectives

The reconstruction of causal networks can be potentially extended to various data types. For example, other causal anchors to infer causal gene networks may include epigenetic markers, copy number variations, and perturbation screens. By targeting tissue-specific or species-specific genes, Findr may also reconstruct cross-tissue or host-pathogen/microbiota causal gene networks. The same analysis may also apply on gene isoforms, proteome, etc, to reconstruct multi-omics causal networks. By considering different distributions of technical variation, the very same causal inference may also infer cell type-specific causal networks from single-cell datasets. Each of those perspectives contains its unique challenges that are worth addressing in the future. However, the statistical and computational frameworks have already been laid down.

## Acknowledgements

Development of Findr was supported by grants from the BBSRC [BB/J004235/1, BB/M020053/1].

